# Seasonality of the estrus cycle in laboratory mice under constant conditions

**DOI:** 10.1101/2024.09.18.613702

**Authors:** Tess M. Reichard, Caitlin H. Miller, Jay Yang, Michael J. Sheehan

## Abstract

Seasonality governs every aspect of life in the natural environment. Controlled laboratory settings are intended to keep animals under a constant set of environmental cues with no seasonality. However, prior research suggests that seasonal variation may exist despite aseasonal lab environments. Here, we examined if seasonal reproductive variation was present in a laboratory mouse strain (C57BL/6J) under standard laboratory housing conditions. We found that female C57BL/6J mice exhibited reproductive seasonality mirroring the outside environment, in a controlled “simulated summer” environment. In the winter and spring, females have longer ovulating phases (proestrus and estrus), compared to the fall. Females similarly experience lengthier complete cycles in the spring, with the most rapid cycling occurring in the fall. Additionally, females spent more time in ovulating phases across seasons than previously reported. Laboratory mice are sensitive to external seasonal changes, despite their local environment being light, temperature, and humidity controlled. This may be due to the detection of an unidentified external cue providing information about external seasonal changes. These findings represent just one example of how seasonality may impact mouse physiology in laboratory settings, emphasizing the need to account for such influences in biomedical research.

---

Seasonality represents a critical selective force that drives diverse morphological, physiological, and behavioral adaptations across all forms of life (1–4). By tracking external cues such as photoperiod, animals can anticipate upcoming shifts in the environment (5–9). For example, mammals often grow thicker coats prior to the winter months to stay warm. Many animals similarly exhibit seasonal patterns of reproduction (7,8,10–14). For instance, the shorter photoperiods in the fall and winter trigger reproductive activity in sheep (i.e. short day breeders) (15–17). Short-day seasonal breeders gestate in the winter and give birth in the spring or summer. Even continuous breeders, such as various primate species, show evidence of seasonal patterns due to resource availability, gestation length, and climate (11). Reproductive seasonality generally ensures that offspring become independent when resources and temperatures are more favorable, and thus have the best chances of survival (18,19).

Whether seasonal or continuous breeders, all mammals experience ovulatory estrus cycles. Female mammalian estrus cycles typically consist of four phases: proestrus, estrus, metestrus, and diestrus (20–22). Females in proestrus are approaching ovulation which occurs during estrus, while the metestrus and diestrus phases occur after egg release when females shed their uterine lining in the absence of successful implantation (21–24). Variation in the length and timing of estrus cycles have been observed across a wide range of mammalian species in response to photoperiod or temperature, both of which are essential cues of seasonal changes (1,15,17,18,25,26).

Interestingly, even in controlled captive environments with abundant food and resources, animals often continue to show seasonal reproductive variation (11,26–29). In laboratory settings, environmental conditions, including photoperiod and temperature variation, have been highly regulated for hundreds of generations for most model organisms (30). Despite these regulated conditions, researchers have detected seasonal hormonal surges in laboratory rats, suggesting the possibility of an innate component to seasonality and/or critical cues in laboratory environments that are not accounted for (31).

House mice are one of the most well-studied organisms, and are widely used in all areas of biomedical research, and abundantly in reproductive research due to their relatively short and consistent 4-5 day estrus cycles (20,32–34). However, in most seasonality studies animals with more pronounced seasonal responses are typically used instead of mice (e.g. hamsters and sheep) (16). In the wild, house mice exhibit seasonal breeding patterns, producing most of their offspring in the spring and the fewest in the fall (35–39). Increased temperature and food availability in the spring allow female mice to consume enough to offset energy lost during pregnancy and lactation. Spring-born pups benefit from a resource-rich spring and summer environments compared to those born in fall or winter (35). When pups are 17-23 days old, they transition from nursing to eating solid food, some of which they forage on their own (39–41). Becoming nutritionally independent from their mother and learning how to forage is imperative for pup survival, so being born when resources are becoming plentiful is highly beneficial (35,40,42). In sum, reproductive seasonality improves the outcomes of both mother and offspring by aligning reproduction with periods of resource abundance.

Under laboratory conditions, house mice tend to breed continuously (30,43). Lab mice experience highly regulated environments in which weather and temperature variation are eliminated, photoperiod is consistent, food is abundant, and there is no predation. The standard light procedure in mouse holding rooms is either a 12:12 or 14:10 light:dark cycle, both of which simulate the longer days of summer (43). Despite the seemingly constant conditions of the lab, some early work observed monthly variation in house mouse estrus cycle length in captivity (27,44,45). Laguchev (1959) demonstrated that mice had a longer overall estrus length in the spring, while Watson & Chaykin (1987) showed that there was more frequent and rapid cycling in the late fall and late spring months. Though this work shows variation in overall estrus cycle length, the variation in the separate phases of the estrus cycle remains unknown. By examining the different phases of the estrus cycle, we can gain insight on the adaptive function of reproductive seasonality in mice, especially under nominally controlled conditions and in the most commonly used laboratory strain. Despite hundreds of generations living in captivity, mice may still have longer or shorter phases of their cycle depending on whether natural environmental conditions would be relatively suitable for pups.

## Methods

The objective of this study was to investigate whether seasonal reproductive variation was present in a commonly used genetically inbred laboratory mouse strain under standard laboratory conditions. To do so, we examined 378 vaginal cytology assays performed on 190 female C57BL/6J mice over the course of two years capturing the lengths of 243 complete estrus cycles and 135 partial estrus cycles consisting of only the ovulating (proestrus and estrus) or quiescent phase (metestrus and diestrus). We initially collected estrus cycle data as part of a study on the effects of estrus and pregnancy state on female mouse responses to social odors (46); we collected additional estrus cycle data after we suspected a seasonal pattern was present.

### Housing

We obtained C57BL/6J mice (C57) (JAX stock #000664) from the Jackson Laboratory and maintained them under conditions approved by the Institutional Animal Care and Use Committee of Cornell University (Protocol #2015-0060). We bred most of our mice (156/190) in our colony from previously ordered Jackson Laboratory mice, while 42/190 were directly obtained from the Jackson Laboratory. Mice were acclimated in our laboratory for at least 2 weeks prior to tracking their estrus cycle to limit the impacts of any stress associated with being transitioned to our laboratory environment from Jackson Laboratory’s breeding facilities. Mice had access to both food (Standard Mouse Chow, Teklad LM-485 mouse/rat sterilizable irradiated diet) and water *ad libitum*. Each cage contained a 5” cardboard mouse house with standard corn cob bedding. We housed virgin, female mice in Allentown Static Rodent Cages. We kept cages in a room with a photoperiod of 14 hours of light and 10 hours of dark (lights on at 10:00 AM, simulating summer light patterns), and consistent temperatures of 67-71°F (median = 69°F, mean = 69.1°F). Humidity in the room ranged from 20.5-67% (median = 34%, mean = 38.9%).

Temperature and humidity were recorded daily with AcuRite 00613 Indoor Humidity and Temperature Monitor placed inside the colony room. Animals were 2-5 months old and were group-housed with 2-5 females per cage. Every female in the cage was sampled six out of seven days of the week, for two to three weeks consecutively.

### Vaginal Cytology

We tracked the estrus cycle via vaginal cytology (Figure 1), one to two hours before the beginning of the 14:10 shifted light:dark cycle (performed from 8:00 AM to 9:00 AM, lights on at 10:00 AM) (20,32) We classified estrus stage based on the relative abundance of cell types on the slide (20,32,47) (Figure 1C). During proestrus, there are primarily small nucleated epithelial cells with the occasional cornified epithelial cells (Figure 1C); estradiol and luteinizing hormone (LH) are the highest during the phase, with elevated levels of FSH as well (20,23,32,47,48). The estrus phase consists of mostly cornified epithelial cells and is marked with a withdrawal of estradiol (Figure 1C). Both proestrus and estrus have lower progesterone levels, but higher prolactin levels compared to metestrus and diestrus (23,48). For this reason, we combined proestrus and estrus into the “ovulating phase” for our analyses throughout the paper (Figure 1A: purple). Metestrus shows cornified epithelial cells that are often surrounded by leukocytes (Figure 1C) (20,32,47). During metestrus, progesterone levels begin to increase with levels peaking during diestrus. Diestrus is marked by the overwhelming presence of leukocytes with a low abundance of other cell types (Figure 1C) (20,23,47,48). Since metestrus and diestrus have high progesterone levels, we combined these two phases together as the “quiescent phase” (Figure 1A: gray) (49). By grouping together stages with similar hormonal shifts, we minimized risk of inaccurate cycle stage determination and limited the bias of the person scoring the images. Proestrus is the preovulatory day while the egg is released during estrus (ovulating phase) (50); metestrus and diestrus result in the animal being sexually unreceptive and quiescent (quiescent phase). As a classification control, a random set of cytology slides (n=159) were selected and blindly classified as ovulating or quiescent. There was a 94% classification match between the blindly scored and previously determined estrus phases, indicating robust scoring procedures (Table S6). A new machine learning tool, Object Detection for Estrous Staging (ODES) (51), was also assessed for its accuracy in classifying estrus stages. ODES demonstrated a 91% match with the blindly scored images and a 90% match with the previously scored images (Table S6). The variability observed in ODES was comparable to that of human scorers. Notably, the machine learning tool struggled to accurately identify leukocytes, which are predominantly present during metestrus and diestrus, leading to some misclassifications as ovulating when they were quiescent (Table S6).

**Figure 1:**
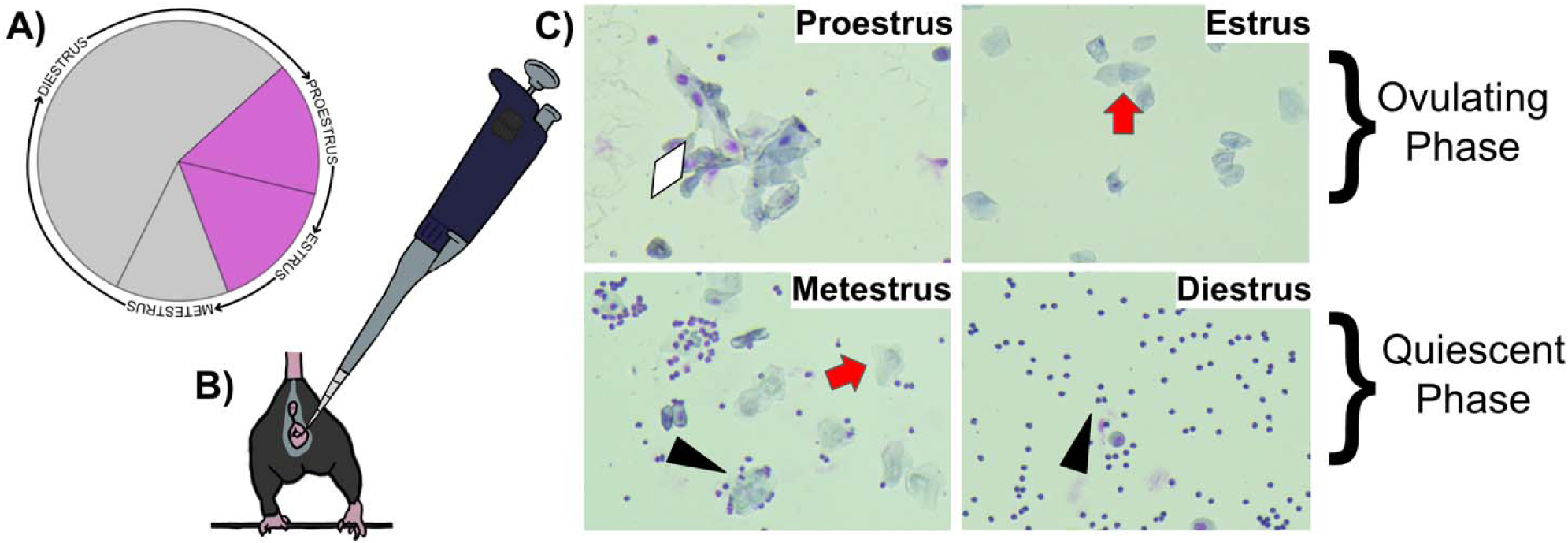
Estrus cycle staging: (**A)** Estrus cycle wheel, showing the relative length of each stage, adapted from Byers et al., 2012 (32). The ovulating phase consists of proestrus and estrus (purple). The quiescent phase consists of metestrus and diestrus (gray). (**B)** Vaginal cytology technique, where a pipette filled with saline solution is placed at the base of the vaginal opening and released gently until saturated with cells. **(C)** The four stages of the estrus cycle within a mouse (proestrus, estrus, metestrus, diestrus). There are three distinct cell types within the estrus cycle, with varying ratios in each stage: cornified epithelial (red arrow), nucleated epithelial (white diamond), and leukocytes (black triangle).

We handled each mouse gently to minimize stress, resting their feet on top of the cage hopper, such that only was elevated during measurement (Figure 1B). We flushed the vagina using 25 µL of Phosphate Buffered Saline (PBS) pH 7.4 (1X), placed at the base of the vaginal opening (Figure 1B)

(20). We repeated the flushing 10 times, or until the solution was cloudy with cells. We then released the solution and smeared it onto a glass slide (stained with JorVet DipQuick Stain Fixative, Stain Solution, and Counter Stain). We washed off any excess stain and transferred slides to a 40X resolution microscope (AmScope T720Q-TP, 40X-1000X). We then photographed the slide (AmScope Microscope Digital Camera, 18 MP Aptina Color CMOS, MU1803) (Figure 1C).

Vaginal cytology was performed repeatedly each day for at least two full cycles for each animal. The start of a new cycle occurred once the mouse transitioned from diestrus to proestrus. Partial cycles included in the data were those where the estrus phase boundaries surrounded both the start and end of the recorded partial cycle. For example, if a mouse was in diestrus one day, in estrus for the subsequent two days, in metestrus for the following day, but there was a data gap after metestrus, only the estrus days (ovulating phase) were included as a partial cycle.

Some estrus cycles contained gaps due to weekends and holidays, which were accounted for in statistical models. There was a large gap in data from the end of December to the end of January, due to an extended break for COVID-19 precautions encouraged by the university (2021-2022) (Figure 3, Figure S2).

### Data Analysis

We analyzed the length of ovulating and quiescent phases using a mixed effect *Cox* model in R (52). Most collected data were complete estrus cycles, but by using this approach, the survival model accounts for right-censored cycles that ended during gaps in measurement (53). Our full model predicted cycle phase length as a function of season (winter, spring, summer, fall), colony room humidity, and colony room temperature (Table S4). We also included random effects of mouse identity and a binary indicator of whether the mice had been bred in our lab or had been acquired from The Jackson Laboratory. Jackson Laboratory origin had significance in the quiescent phase only, so it was included as a random effect in our final model (Table S4). We also included the number of females in each cage in our initial model, but removed this variable from our final model because this factor did not have a significant effect (54). We included average colony room temperature and humidity across each cycle as a fixed effect in our model since season predicted each of these variables, though these were not significant predictors of cycle length (Tables S2, S3, S4). We performed analyses using continuous and categorical measures of time by examining the effect of the day of the year (continuous) and the four astronomical seasons (categorical). Some cycles spanned multiple seasons—in this case we classified them as belonging to the more represented season or randomly if it was evenly split.

Additionally, we built a mixed effects model using the glmmTMB package in R comparing the ovulating phase percentage across seasons (55) (Table 2).

## Results

### **(a)** Reproductive Seasonality

Proestrus/estrus (ovulating phase) length differed significantly across seasons (Figure 2A, Figure 3, Table 1). The duration of the ovulating phase was significantly longer during the spring and winter compared with the fall (*p*=0.003, *Z*=-2.97, OR=0.57 and *p*=0.035, Z=-2.1, OR=0.66, respectively; Figure 2A, Table 1). Metestrus/diestrus (quiescent phase) length was shortest in the fall as compared to spring and summer (spring: *p*<0.0001, *Z*=-3.9, OR=0.46; summer: *p*=0.00021, *Z*=-3.7, OR=0.41; Figure 2B, Table 1). An odds ratio (OR) less than one means that it is less likely to end, signifying that season is longer in duration compared to the reference season. Additionally, the quiescent phases during spring and summer were longer as compared to during winter (spring: *p*=0.027, *Z*=-2.21, OR=0.55; summer: *p*=0.028, *Z*=-2.19, OR=0.49; Figure 2B, Table 1).

**Figure 2:**
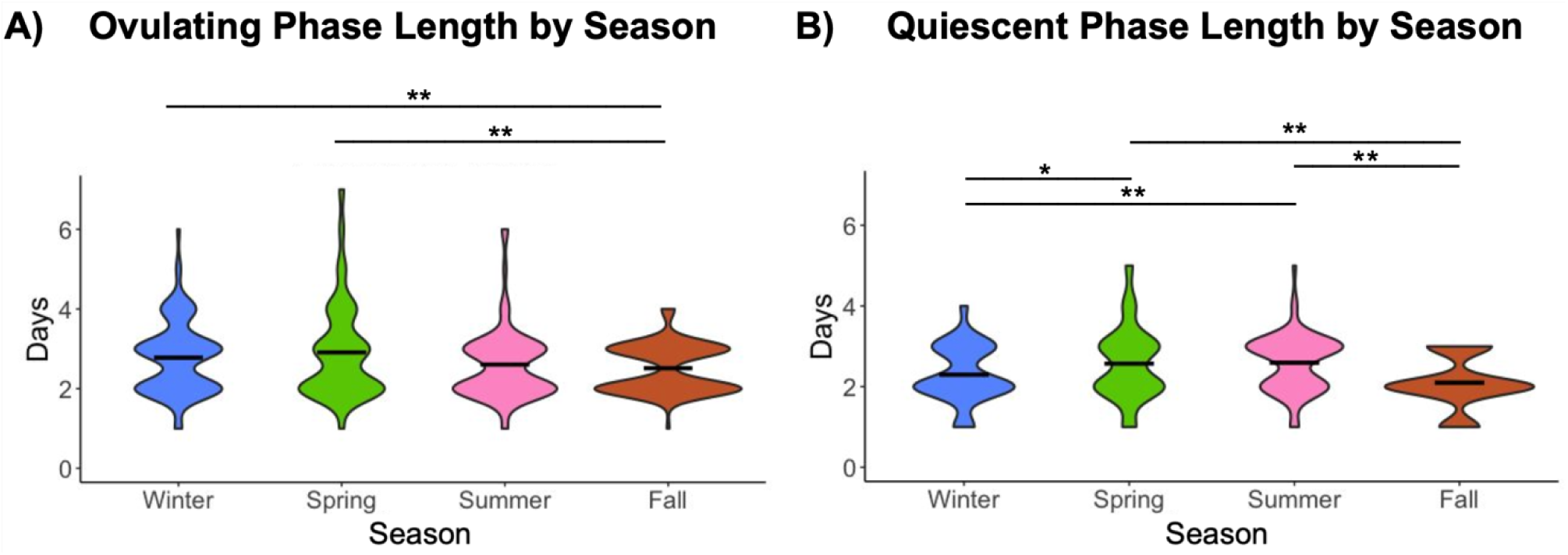
Estrus cycle phase length differences across seasons. Depicted above is a violin plot, showing the density and spread of data points across all seasons. **(A)** Ovulating phase includes proestrus and estrus, **(B)** quiescent phase consists of metestrus and diestrus. Ovulating phase number per season: n=96 for winter, n=46 for spring, n=71 for summer, n=111 for fall. Quiescent phase number per season: n=84 for winter, n=49 for spring, n=64 for summer, n=102 for fall. The black bars indicate the mean number of days per season. Ovulating phase mean days: winter=2.83, spring=3.02, summer=2.60, fall=2.51. Quiescent phase mean days: winter=2.30, spring=2.57, summer=2.59, fall=2.09. Larger bulges within each violin signify a larger density of data (ovulating phase median days: winter=3, spring=3, summer=2, fall=2; quiescent phase median days: winter=2, spring=2, summer=3, fall=2). Lines above the plot show significance between two seasons using a survival model, where ** indicates p < 0.01 and * indicates p < 0.05.

**Figure 3:**
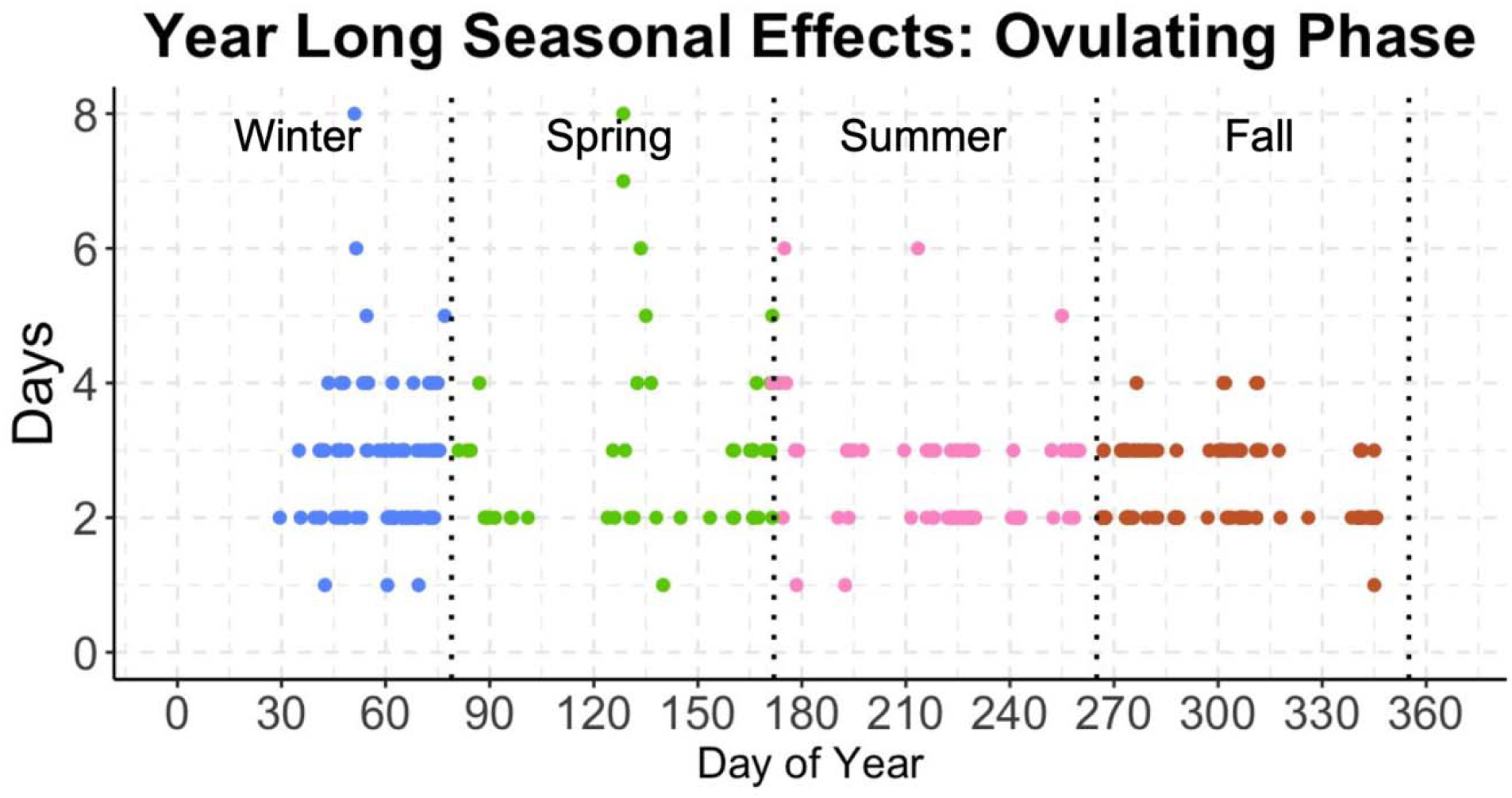
Julian plot showing the distribution of data points throughout the year. Each data point is the midpoint of a female’s ovulating phase length, plotted by season (n=324). The ovulating phase plotted over the year separated by season. The black dotted vertical lines indicate seasonal boundaries. The color of the data points shows the season in which the ovulating phase was recorded, and the quadrant each point lies in shows the season in which the pup would be born. Day 0 represents January 1, whereas Day 365 represents December 31.

**Table 1:**
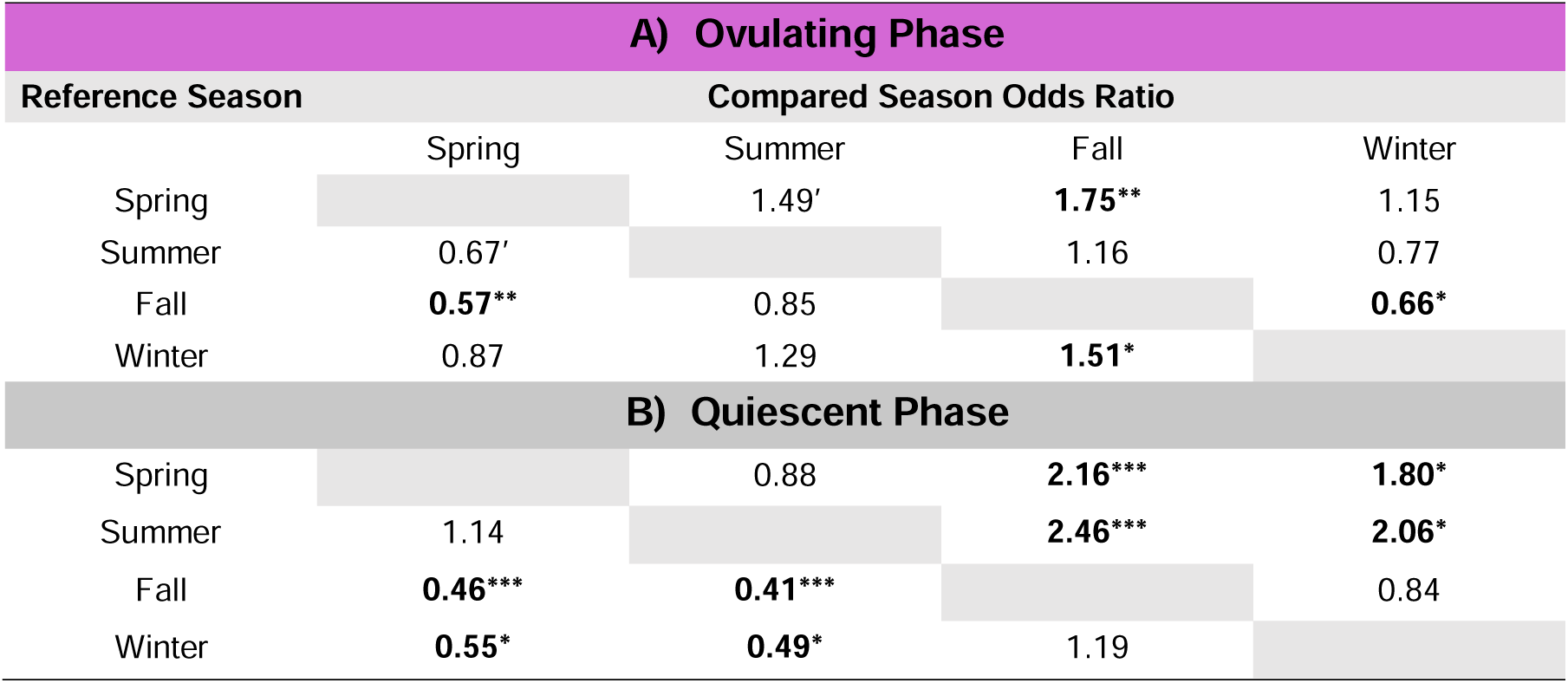
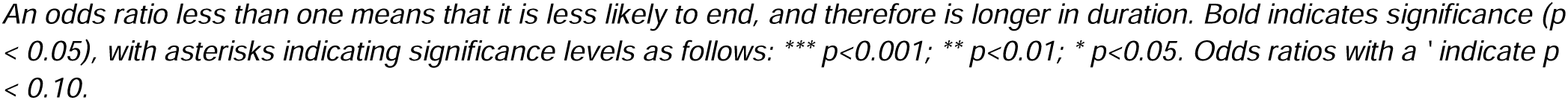
The odds ratio of estrus cycle phases ending **(A)** Ovulating Phase **(B)** Quiescent Phase.

| A) Ovulating Phase |  |  |  |  |
| --- | --- | --- | --- | --- |
| Reference Season | Compared Season Odds Ratio |  |  |  |
|  | Spring | Summer | Fall | Winter |
| Spring |  | 1.49' | **1.75\*\*** | 1.15 |
| Summer | 0.67' |  | 1.16 | 0.77 |
| Fall | **0.57\*\*** | 0.85 |  | **0.66\*** |
| Winter | 0.87 | 1.29 | **1.51\*** |  |
| B) Quiescent Phase |  |  |  |  |
|  | Spring | Summer | Fall | Winter |
| Spring |  | 0.88 | **2.16\*\*\*** | **1.80\*** |
| Summer | 1.14 |  | **2.46\*\*\*** | **2.06\*** |
| Fall | **0.46\*\*\*** | **0.41\*\*\*** |  | 0.84 |
| Winter | **0.55\*** | **0.49\*** | 1.19 |  |

An odds ratio less than one means that it is less likely to end, and therefore is longer in duration. Bold indicates significance ( $p < 0.05$ ), with asterisks indicating significance levels as follows: \*\*\* $p < 0.001$ ; \*\* $p < 0.01$ ; \* $p < 0.05$ . Odds ratios with a ' indicate $p < 0.10$ .

**Table 2:**
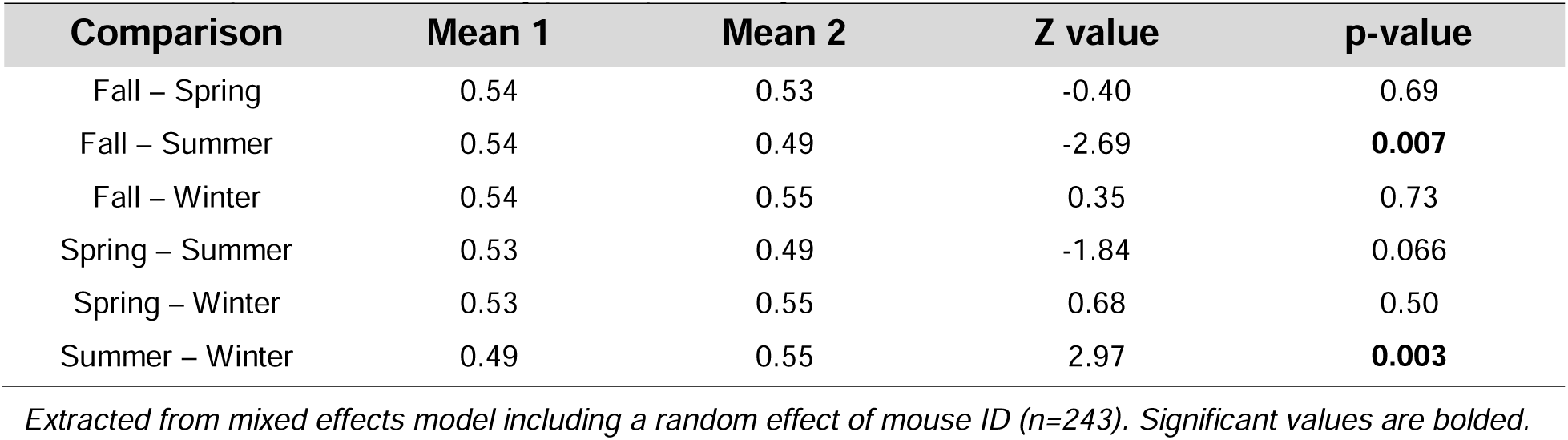
Comparisons of ovulating phase percentage across seasons.

| Comparison | Mean 1 | Mean 2 | Z value | p-value |
| --- | --- | --- | --- | --- |
| Fall – Spring | 0.54 | 0.53 | -0.40 | 0.69 |
| Fall – Summer | 0.54 | 0.49 | -2.69 | **0.007** |
| Fall – Winter | 0.54 | 0.55 | 0.35 | 0.73 |
| Spring – Summer | 0.53 | 0.49 | -1.84 | 0.066 |
| Spring – Winter | 0.53 | 0.55 | 0.68 | 0.50 |
| Summer – Winter | 0.49 | 0.55 | 2.97 | **0.003** |

*Extracted from mixed effects model including a random effect of mouse ID (n=243). Significant values are bolded.*

Ovulating phase length was graphed as a Julian plot to visualize the day each data point was collected during the year (Figure 3). Ovulating phases in winter and spring showed the widest length distribution, while fall had the least, ranging from 1 to 4 days (Figure 2, Figure 3). Despite consistent sampling across fall, no ovulating phases in the fall were observed to be longer than 4 days (Figure 2, Figure 3).

In contrast, 4.2% of winter and 10.9% of spring ovulating phases were longer than 5 days, with phases as long as 8 days observed in both seasons (Figure 3). The length of the entire estrus cycle (i.e. “full cycle”, including all 4 phases) also significantly differed across all seasons in relation to the fall (Figure S1, Table S1). Females in spring experienced longer full cycles compared to the fall cycle length (spring: *p*=0.0026, *Z*=-3.01, OR=.51; Table S1).

### **(b)** Estrus Cycle Proportion

It has been frequently reported that metestrus and diestrus have the longest duration during the estrus cycle, while proestrus and estrus are relatively shorter phases (Figure 4A). We found that the proportion of time females spent in the ovulating phase was greater than prior reports (20,30,32,33,56) (Figure 4). This ratio also varied across the seasons, rising from 45:55 (quiescent:ovulating) in winter to 46:54 in spring, reaching a peak of 50:50 in summer and declining to 48:52 in fall (Figure 4B). Fall and winter have a higher ovulating phase percentage compared with summer (Figure 4C, Table 2). The higher ovulating percentage in fall may be due to fall having the shortest overall estrus cycle length (Figure S1, Table S1). Across all seasons, the ovulating and quiescent phase are similar in duration (Figure 4B), with the ovulating phase constituting most of the estrus cycle. In addition, the length of the estrus cycles that we measured exceeded the previously published range of 4-5 days; the range in our animals was 3-11 days (20,57) (Figure 4, Figure S2). The one-day gaps observed in the unusually long cycles were consistently between two instances of a known phase, such as the quiescent phase. For example, if the preceding day before the gap was in the quiescent phase and the succeeding day was also in the quiescent phase, it suggests continuity within the same phase. This pattern indicates that it is unlikely the missing day involved a complete switch to the ovulating phase. However, while we cannot entirely rule out the possibility of missing a phase transition, we believe it is improbable.

**Figure 4:**
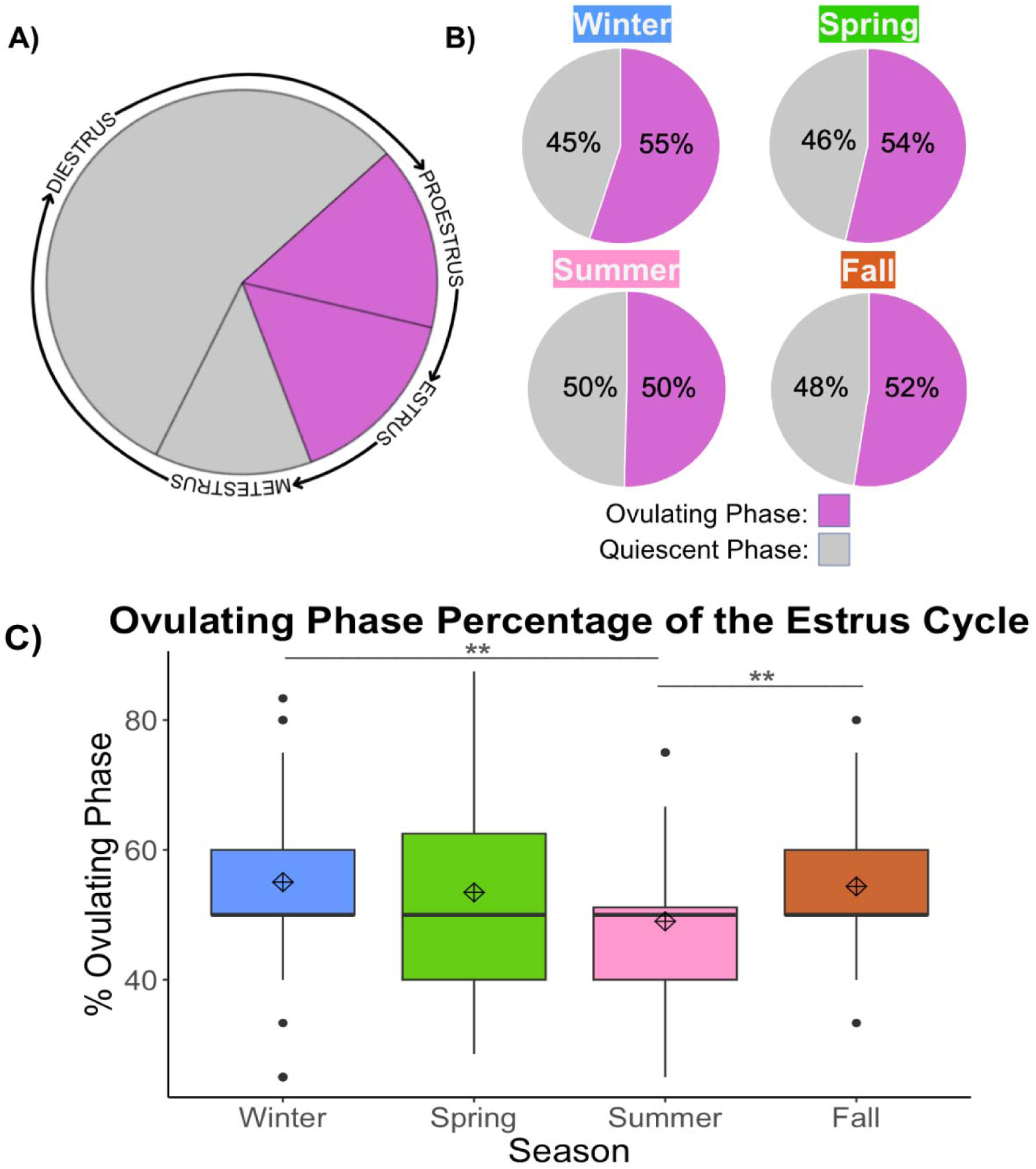
Estrus cycle proportion. (**A)** Estrus cycle wheel, showing the relative length of each stage, adapted from Byers et al., 2012 (32). **(B)** Observed proportion of cycling, where proestrus and estrus are grouped together, and metestrus and diestrus are grouped together. **(C)** The percent of ovulating phase within the estrus cycle by season. Cycle number per season: n=70 for winter, n=37 for spring, n=60 for summer, n=76 for fall. Black lines indicate the median; diamond with cross indicates the mean. Mean percentage: winter=55.04, spring=53.45, summer=49, fall=54.37. Median percentage: winter=50, spring=50, summer=50, fall=50. Lines above the plot show significance between two seasons using a linear mixed effects model, where ** indicates p < 0.01.

## Discussion

Even within a controlled laboratory environment resembling “eternal summer,” we observed seasonal reproductive effects on estrus cycle length in inbred C57BL/6J female mice. Ovulating phases in winter and spring were longer in duration than those in fall (Figure 2A, Table 1). Similarly, complete estrus cycles were longest in spring, with the shortest cycle duration in fall (Figure S1, Table S1). We additionally found that the ovulating and quiescent phase were approximately evenly represented (Figure 4), in contrast to previous studies on estrus cycle durations. C57 mice have been bred in the lab for hundreds of generations (though see X for information about the genetic stability program for common strains at the Jackson Lab) and display distinct behavioral phenotypes compared to wild mice, including a reduction in aggression and an increase in disease susceptibility (43,58–62). The inbreeding and stable lab setting eliminate genetic and largely eliminate environmental variation, respectively, yet mice still demonstrate reproductive variation.

These results have implications in all research fields utilizing mice as a model for reproductive physiology, especially those focused on intervention. Failure to account for seasonal fluctuations in these analyses risk overlooking seasonal physiological differences. Biomedical research often dismisses external variables in experiments (63–68); however, our research reveals that mice alter their physiology in response to seasonal changes. Although we only report results on estrus cycle lengths, other physiological and behavioral processes are likely also affected by seasonal changes, as the estrus cycle directly reflects hormonal changes (9,23). These female mice are not receiving any apparent cues from their controlled environment, yet still determine the external seasonality and adapt their estrus cycle accordingly.

What environmental cues might the animals have used to drive their seasonal fluctuations in reproductive physiology? It is conceivable, that the brief transportation from the Jackson Laboratory to Cornell University might have provided sufficient cues to the mice for interpreting season, allowing them to entrain longer term circannual rhythms. When C57 mice were ordered from the Jackson Laboratory (42/190), they were shipped in a breathable transportation box from a distributor to Cornell University.

The short period of transportation temporarily exposed the mice to outside weather conditions, such as temperature, humidity, and precipitation. However, most of our study animals were bred in our lab (148/190). These animals also showed seasonal variation in cycle length and animals’ site of origin did not predict cycle length. We performed analyses using only Jackson Laboratory-bred or only colony-bred mice, but the same seasonal patterns were apparent for both groups. Thus, we do not believe that transport from Jackson Laboratory accounts for the observed seasonal variation. We also found that humidity and temperature in our mouse colony room did not predict variation among estrus cycle length (Tables S2, S3, S4).

An alternative cue of seasonality may be subtle differences in experimenter and laboratory variables that changed over the seasons. For example, differences in animal physiology have been previously documented in response to the sex of the researcher, where male researchers elicit a stress response in mice (69,70). Furthermore, results obtained from one laboratory often cannot be replicated in another (63,65,66,68,71–73). Researchers do not know why there are discrepancies in results but know there are some environmental factors unique to each lab which affect the results, called the “unknown unknowns” (65). Although laboratory conditions are relatively standardized, there are still differences in experimental outcomes, in which the identity of these laboratory artifacts have not been determined.

For this reason, perhaps mice in some labs might experience seasonality and others will not, due to unknown environmental or experimenter cues.

It is therefore likely that mice use some unknown cue in the laboratory environment to entrain a circannual rhythm of which we are unaware. Researchers may be unknowingly bringing in a cue from outside on their clothes or skin, which could be sufficient for mice to interpret the seasonality. As a result, the estrus cycles of mice should be tracked in other lab colonies that have different seasonality than Ithaca, NY, including regions with constant conditions and inverse seasonal patterns in the southern hemisphere. For example, regions near the equator undergo minimal fluctuations in both photoperiod and temperature, with predominant shifts occurring in precipitation, leading to distinct wet and dry seasons (74). Tracking the estrus cycle of laboratory females in these regions would determine the presence of seasonal patterns and could help pinpoint the specific cues mice rely on. Additionally, researchers could change the overt seasonal cues in a mouse holding facility. For example, one could rear mice under a winter light cycle (shorter daylight, longer darkness) in summer months, and observe if seasonal trends still exist (22). Alternatively, increasing precautions when entering the rearing facility—such as removing outside clothes, showering, and changing into clean clothes—could potentially eliminate the observed seasonal variation; standardization of this protocol may mitigate the observed seasonal effects.

The changes we observe in estrus cycles are consistent with predicted changes that one would expect to observe in free-living temperate rodent populations. Tradeoffs are necessary for survival since every action incurs some kind of cost to an animal, especially reproductive processes. Reproduction is particularly energetically costly, in which females prepare for it by increasing their body fat reserves, foraging, or hoarding food (75–77). Schneider et al. (2017) showed that female hamsters fed *ad libitum* abandoned their natural food hoarding behavior (75,78), but when females became food-restricted, they began to hoard for 75% of their estrus cycle. These hamsters redirected metabolic energy that could have gone towards reproduction to hoarding food (75). Therefore, as environmental factors become constraining to their metabolic balance, females make tradeoffs, incur reproductive opportunity costs, and adjust their behavior to optimize their anticipated external environment (75,79). Due to the absence of environmental constraints and selective pressures in the laboratory setting, female mice may experience a prolonged ovulating phase. This could explain the more equal proportion of quiescent:ovulating seen in the C57 mice in our study (Figure 4). The increased ovulating phase duration might be because there are no pressures to control the success and growth in offspring. This increase in duration may also be more energetically expensive, so females may have to cope with a longer overall cycle length to recover. This could be responsible for the longer estrus cycle lengths seen in the spring.

An alternative reason for longer estrus cycle lengths could be due to the domestication of the house mouse, which has led to various genetic, physiological, and morphological changes that differ from wild populations (43,80). Animals generally become less aggressive, more explorative, and sexually mature at a younger age (80). Additionally, prior to attempts to stabilize the genetic makeup of mouse lines (), researchers may have inadvertently selected for rapid breeding by increasing the window of fertility in lab mice, and thus increasing the chances of a successful copulation. Female behavior and physiology have likely evolved in response to domestication in house mice, to maximize breeding efficiency in laboratory settings (59,62).

Mice in our study exhibited longer ovulating phases in spring and winter, which would potentially optimize offspring survival and maturity during seasons with abundant resources and favorable temperatures in the wild. A similar pattern is observed in seasonally breeding deer, where mating occurs in the fall, females are pregnant in the winter, and fawns are birthed in late spring and summer; though if deer are not bred in the fall, they will continue cycling through March (18,81,82). Wild populations of house mice are seasonal breeders in which offspring dispersal occurs about seven weeks after the initial conception (35). The increased length in ovulating phase observed in winter and spring would increase the probability of females getting pregnant and are consistent with the breeding season in wild house mouse populations (35–37,40,83). For this reason, fall may have the shortest estrus cycle due to winter being the following season (Figures 2, S1; Table S1). With temperature drops and restricted food resources, it is likely advantageous for wild mice to minimize the instances of pregnancy investment during this period. So, while mice living under natural conditions might halt estrus altogether in the late fall, under the less energetically constrained conditions of the lab, this circannual rhythm appears to be dampened but still present.

The main limitation with our study is the data gaps caused by interrupted sampling. While our survival model accounted for these gaps, complete estrus cycle data would have made this unnecessary, though it requires daily tracking, which is not always practical. We also combined estrus stages which reduced scorer bias and unified similar hormonal shifts; additionally, proestrus is very short lasting, so it was often missed during our daily assays (20,32,48). Regardless, our findings convey that even in seemingly environmentally controlled laboratory settings mice are able to pick up on external cues relating to outside environment.

Seasonal changes in cycle length may have downstream consequences for offspring survival. A prior lab study found that house mice born in winter had the longest lifespan, while those born in summer had the shortest, with similar outcomes observed in rats and fruit flies (84). Future work should study reproductive patterns alongside survival outcomes to provide insights into how organisms might strategically time their reproductive efforts to align with seasons that offer optimal conditions for offspring survival. Since laboratory survival of house mice coincides with wild population survival (35,39,40,42,84), we see that their reproductive strategy is maintained across both environments with no apparent cause.

## Conclusions

C57 mice exhibit seasonal effects in their reproductive cycle in the absence of key seasonal cues, such as photoperiod, temperature, humidity, or food availability. This suggests there is an unknown cue or physiological mechanism controlling seasonal reproductive variation when no apparent environmental information is available. Though this mechanism of seasonality is unknown, mice may gather environmental cues within a laboratory environment which go undetected by humans. These results suggest that house mice have a photoperiod-independent mechanism for tracking seasons or can use cues brought in by researchers to seasonally adjust. Researchers, especially with studies involving mice, rarely mention the time of year it was performed. Critically, results may not be replicable if the experiment were performed at a different time of the year, since seasonality is observed in controlled mouse colonies.

## List of abbreviations

C57: C57BL/6J
JAX: The Jackson Laboratory
OR: Odds Ratio

## Declarations

### Ethics approval and consent to participate

All experimental procedures were approved by the Cornell University Institutional Animal Care and Use committee (IACUC: Protocol #2015-0060) and followed the NIH Guide for Care and Use of Animals.

### Consent for publication

All authors read and approved the final manuscript for publication.

### Availability of data and materials

All statistical code and outputs are included in **Additional File 3**. Metadata is included in **Additional File 1.** Estrus cycle reclassification is included in **Additional File 2.**

## Authors’ contributions

Conceptualization: CHM. Study design: CHM, TMR. Methodology design: CHM, TMR. Data curation and code: TMR, JY, CHM. Data analysis: TMR, CHM. Figure creation: TMR. Writing—original draft: TMR, CHM. Writing—review and editing: TMR, CHM, JY, MJS. Funding acquisition: MJS.

## Supporting information

Additional File 3 - R code for analysis

## Acknowledgements

We thank Matthew N. Zipple and Caleb C. Vogt for helpful discussions, statistical consultation, and manuscript comments; Melanie Colvin and Jeremy Cusker for technical assistance.

