## Supplemental Tables and Figures for "Seasonality of the estrus cycle in laboratory mice under constant conditions"

**Supplementary Tables**

***Table S1:*** *The odds ratio of the full estrus cycle ending*

| **Full Cycle** | | | | |
| --- | --- | --- | --- | --- |
| **Reference Season** | **Compared Season Odds Ratio** | | | |
|  | Spring | Summer | Fall | Winter |
| Spring |  | 1.14 | **1.96**** | 1.34 |
| Summer | 0.87 |  | 1.71’ | 1.17 |
| Fall | **0.51**** | 0.58’ |  | 0.68’ |
| Winter | 0.75 | 0.85 | 1.46’ |  |

*The odds ratio of the full estrus cycle ending. A number less than one means that it is less likely to end, and therefore is longer in duration. Bolded numbers with ** p < 0.01, ‘ p < 0.10*

***Table S2:*** *Comparisons of colony room humidity across seasons*

| **Comparison** | **Mean 1** | **Mean 2** | **t value** | **p-value** |
| --- | --- | --- | --- | --- |
| Fall – Spring | 38.30 | 39.89 | 0.77 | 0.44 |
| Fall – Summer | 38.30 | 59.05 | 9.87 | **<0.0001** |
| Fall – Winter | 38.30 | 24.54 | -7.22 | **<0.0001** |
| Spring – Summer | 39.89 | 59.05 | 8.51 | **<0.0001** |
| Spring – Winter | 39.89 | 24.54 | -7.42 | **<0.0001** |
| Summer – Winter | 59.05 | 24.55 | -16.27 | **<0.0001** |

*Extracted from linear model (humidity ~ season); significant values are bolded*

***Table S3:*** Comparisons of colony room temperature across seasons

| **Comparison** | **Mean 1** | **Mean 2** | **t value** | **p-value** |
| --- | --- | --- | --- | --- |
| Fall – Spring | 69.13 | 68.7 | -4.04 | **<0.0001** |
| Fall – Summer | 69.13 | 68.38 | -6.88 | **<0.0001** |
| Fall – Winter | 69.13 | 70.01 | 8.91 | **<0.0001** |
| Spring – Summer | 68.71 | 69.14 | -2.75 | **.007** |
| Spring – Winter | 68.71 | 68.39 | 12.22 | **<0.0001** |
| Summer – Winter | 68.39 | 70.02 | 14.83 | **<0.0001** |

*Extracted from linear model (temperature ~ season); significant values are bolded*

***Table S4:*** Output from full model used in manuscript using fall as the reference season

| **Compared Season** | **coef** | **exp(coef)** | **se(coef)** | **Z value** | **p-value** |
| --- | --- | --- | --- | --- | --- |
| A) **Ovulating Phase** | | | | | |
| Spring | -0.56 | 0.57 | 0.19 | -2.97 | **0.003** |
| Summer | -0.16 | 0.85 | 0.20 | -0.77 | 0.44 |
| Winter | -0.42 | 0.66 | 0.20 | -2.11 | **0.04** |
| Humidity | 0.002 | 1.00 | 0.007 | 0.35 | 0.83 |
| Temperature | 0.048 | 1.05 | 0.13 | 0.36 | 0.74 |
| B) **Quiescent Phase** | | | | | |
| Spring | -0.77 | 0.46 | 0.20 | -3.90 | **0.0001** |
| Summer | -0.90 | 0.41 | 0.24 | -3.70 | **0.0002** |
| Winter | -0.18 | 0.84 | 0.19 | -0.94 | 0.35 |
| Humidity | 0.01 | 1.01 | 0.007 | 1.44 | 0.15 |
| Temperature | -0.021 | 0.98 | 0.14 | -0.15 | 0.88 |
| C) **Quiescent Phase** | | | | | |
| Spring | -0.67 | 0.51 | 0.22 | -3.01 | **0.003** |
| Summer | -0.54 | 0.58 | 0.28 | -1.89 | 0.059 |
| Winter | -0.38 | 0.68 | 0.22 | -1.76 | 0.079 |
| Humidity | 0.008 | 1.01 | 0.009 | 0.94 | 0.35 |
| Temperature | 0.096 | 1.10 | 0.15 | 0.65 | 0.51 |

*Extracted from Cox mixed model: Surv(cyclelength, end_cyclelength) ~ season+humidity+temperature+(1|mouseID)+(1|jax)*

*Significant values are bolded; humidity and temperature do not show significance*

**Supplementary Figures**

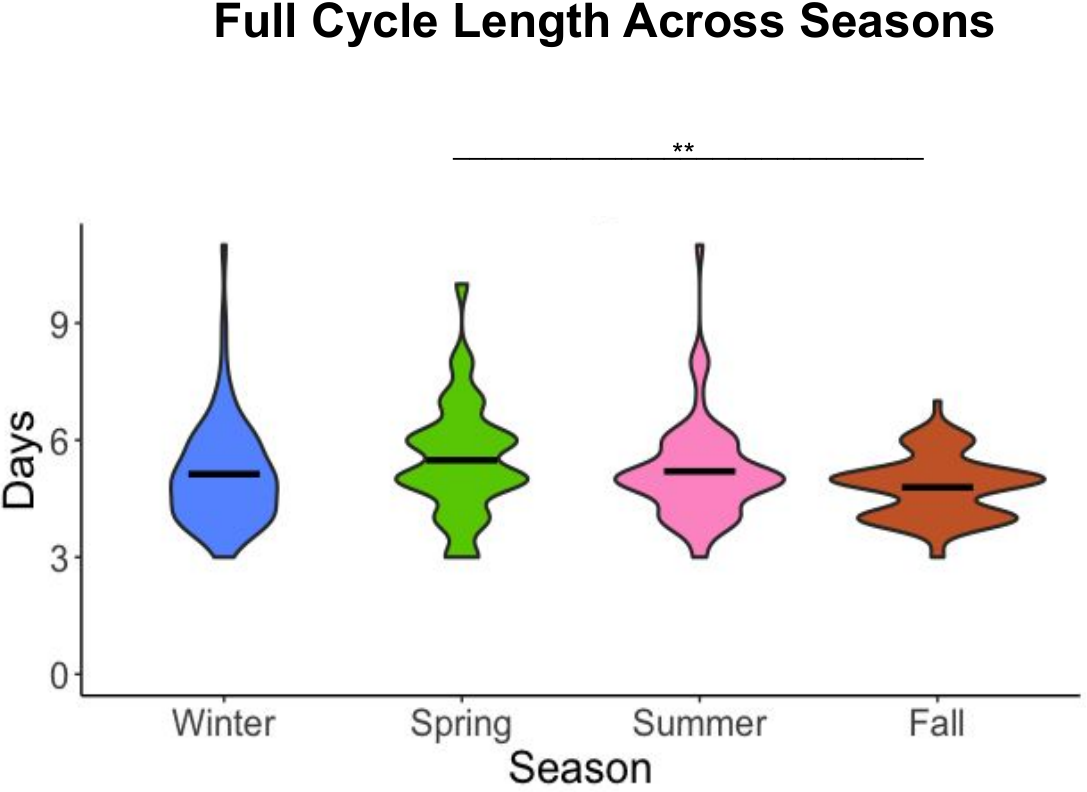

**Figure S1: Full cycle length across seasons.** Larger bulges within each violin signify a larger density of data. The black bars indicate the mean number of days per season. Mean days: winter=5.13, spring=5.49, summer=5.2, fall=4.79. Median days: winter=5, spring=5, summer=5, fall=5. Cycle number per season: n=70 for winter, n=37 for spring, n=60 for summer, n=76 for fall. Lines above the plot show significance between two seasons using a survival model, where ** indicates p < 0.01. Full cycles in spring are longer than full cycles in fall (Table S1).

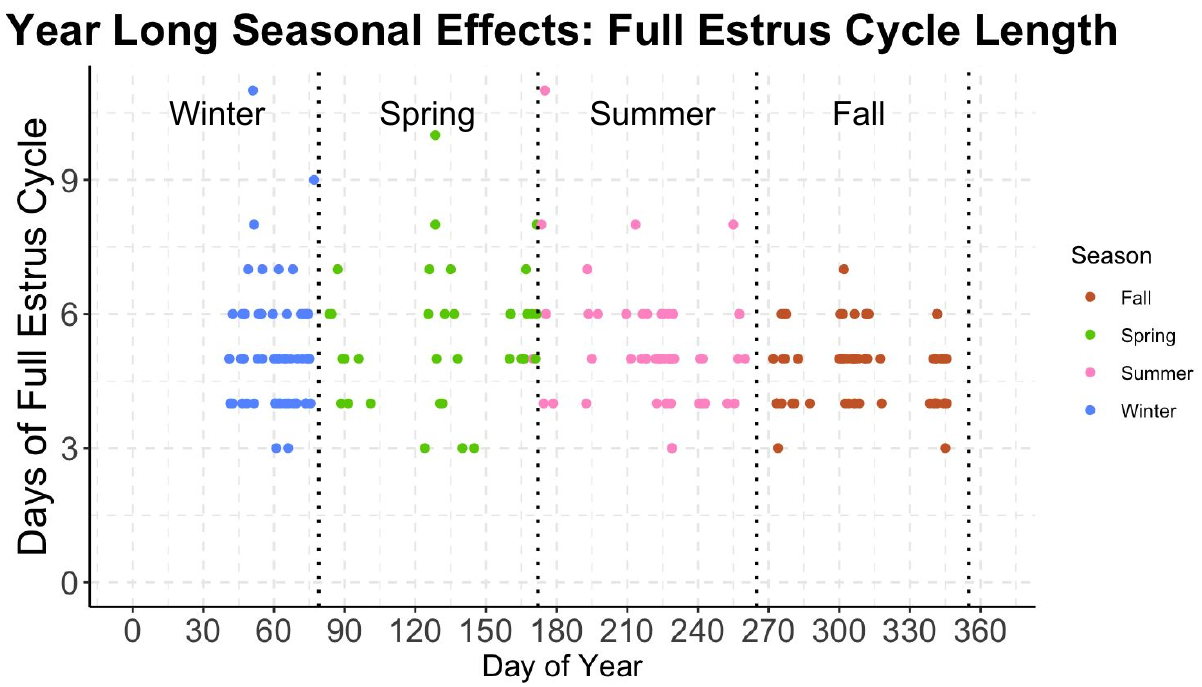

**Figure S2: Julian plots for full estrus cycle length throughout the year.** Each data point is the midpoint of a female’s complete estrus cycles, plotted by season (n=243). The black dotted vertical lines indicate seasonal boundaries. The color of the data points shows the season in which the ovulating phase was recorded, and the quadrants represent seasonal separations. Day 0 represents January 1, whereas Day 365 represents December 31.

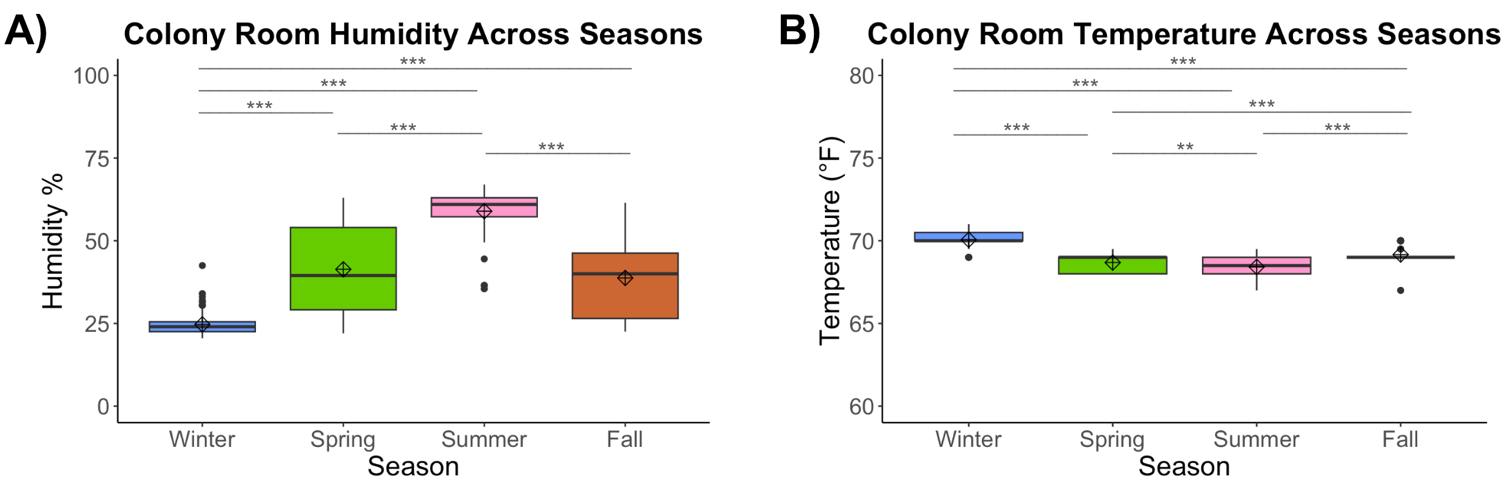

**Figure S3: Temperature and Humidity in Colony Room. (A)** Humidity percentage in the mouse colony room across seasons. Mean humidity percentage: winter=24.5, spring=38.9, summer=59.1, fall=38.3. Median humidity percentage: winter=24, spring=35, summer=61, fall=38.5.  **(B)** Temperature in the mouse colony room across seasons. Mean temperature (°F): winter=70, spring=68.7, summer=68.4, fall=69.1. Median temperature (°F): winter=70, spring=69, summer=68.5, fall=69. Black lines indicate the median; diamond with cross indicates the mean. Lines above the plot show significance between two seasons using a linear model, ** p<0.01, *** p < 0.0001; season predicted colony room humidity and temperature (Tables S2 and S3). Humidity and temperature did not predict the length of the ovulating phase, quiescent phase, or full estrus cycle (Table S4).

**Supplementary Files**

**Additional File 1**

**Table S5: Metadata**

Metadata information for all mice.

**Additional File 2**

**Table S6: Estrus cycle classification match**

A spreadsheet showing the previously classified estrus stages with blindly re-classified estrus stages, demonstrating a 94% match. Previously classified and blindly classified estrus stages were also compared to classifications performed by Object Detection for Estrous Staging (ODES) model (51), demonstrating a 89.9% match and 91.2% match, respectively.

**Additional File 3**

R script showing both data analyses used in the paper and preliminary data analyses. Full statistical outputs for all analyses and figures.
